# A massively parallel strategy for STR marker development, capture, and genotyping

**DOI:** 10.1101/063727

**Authors:** Logan Kistler, Stephen M. Johnson, Mitchell T. Irwin, Edward E. Louis, Aakrosh Ratan, George H. Perry

**Affiliations:** Departments of Anthropology and Biology, Pennsylvania State University, University Park, PA 16802 USA; School of Life Sciences, University of Warwick, Coventry, CV4 7AL United Kingdom; Department of Anthropology, Northern Illinois University, DeKalb, IL 60115 USA; Center for Conservation and Research, Omaha’s Henry Doorly Zoo and Aquarium, Omaha, NE 68107 USA; Department of Public Health Sciences and Center for Public Health Genomics, University of Virginia, Charlottesville 22908, VA, USA

## Abstract

Short tandem repeat (STRs or microsatellites) variants, are highly polymorphic markers that facilitate powerful, high-precision population genetic analyses. STRs are especially valuable in conservation and ecological genetic research, yielding detailed information on population structure and short-term demographic flux. However, STR marker development and analysis by conventional PCR-based methods imposes a workflow bottleneck and is suboptimal for noninvasive sampling strategies such as fecal DNA recovery. While massively parallel sequencing has not previously been leveraged for scalable, efficient STR recovery, here we present a pipeline for developing STR markers directly from high-throughput shotgun sequencing data without requiring a reference genome assembly, and a methodological approach for highly parallel recovery of enriched STR loci. We first employed our approach to design and capture a panel of 5,000 STR loci from a test group of diademed sifakas (*Propithecus diadema*, n=3), endangered Malagasy rainforest lemurs, and we report extremely efficient recovery of targeted loci—97.3-99.6% of STRs characterized with ≥10x non-redundant coverage. Second, we tested our STR capture strategy on a *P. diadema* fecal DNA preparation, and report robust initial results and methodological suggestions for future implementations. In addition to STR targets, this approach also generates large, genome-wide single nucleotide polymorphism (SNP) panels from regions flanking the STR loci. Our method provides a cost-effective and highly scalable solution for rapid recovery of large STR and SNP datasets in any species without need for a reference genome, and can be used even with suboptimal DNA, which is more easily acquired in conservation and ecological genetic studies.

**Data Deposition:** Raw sequencing data are available under Study Accession numbers SRP073167 (genomic shotgun data for Oberon and Tatiana) and SRP076225 (targeted re-sequencing data) from the NCBI Sequence Read Archive. BaitSTR software is available at Github (core BaitSTR programs: https://github.com/aakrosh/BaitSTR; BaitSTR_type.pl companion script for genotyping and block manipulation: https://github.com/lkistler/BaitSTR_type).

## Introduction

Short tandem repeats (STRs; microsatellites) are highly variable genetic markers useful for a wide variety of applications in population genetics. Germline STR mutability is driven by DNA polymerase slippage over tandem repeat regions of short sequence motifs, yielding highly variable copy number repeats (Ellegren 2004; Fan and Chu 2007; Willems et al. 2014). Because of their high-resolution variability at the population level, even across fine temporal scales, STRs have been used to study short-term population demographics and gene flow, identify conservation units, quantify population health in endangered species, and study the genetic basis of behavior in natural populations (Veeramah and Hammer 2014; Hoban et al. 2013; Quéméré et al. 2012). Traditional methods for STR genotyping rely on PCR amplification of genomic targets containing STRs, followed by copy number inference from amplicon fragment size estimated through electrophoresis (Guichoux et al. 2011). For high-quality DNA samples, these methods are proven, accurate, and inexpensive at project scales, but they introduce an unavoidable workflow bottleneck and restrict the scale of analysis to dozens of markers. Due to high exogenous DNA content, the presence of PCR inhibitors, and allelic dropout issues, STR genotyping using these methods with lower-quality, non-invasive samples (e.g. DNA extracted from feces) is more challenging and even less efficient (Arandjelovic et al. 2009; Buchan et al. 2005; Mckelvey and Schwartz 2004).

Single nucleotide polymorphism (SNP)-based analyses have come to dominate the genomic era of DNA sequencing and genotyping. STR genotyping has, for the most part, not yet utilized the full potential of massively parallel sequencing for efficient population-scale analyses during this same era. While a wide variety of strategies for population genomic-scale SNP data collection have been developed, even for ancient DNA (Fu et al. 2013; Haak et al. 2015; Carpenter et al. 2013) and difficult sources such as fecal samples (Perry et al. 2010; Chiou and Bergey 2015; Snyder-Mackler et al. 2016), similar methods have not yet been applied widely to STRs. However, because STRs evolve at orders-of-magnitude greater rates than SNPs, efficient massively parallel sequencing-based tools for their recovery and analysis at the population level would be extremely useful for studies requiring fine-scale geographic or recent temporal resolution (Perry 2014).

A handful of strategies have emerged for applying the genomic sequencing toolkit to components of STR genotyping and analysis. STR discovery has been facilitated by shotgun sequencing with the 454 platform (no longer commercially available), yielding markers that were then analyzed using conventional PCR-based genotyping (Schoebel et al. 2013). Inversely, amplicon sequencing of multiplex PCR-amplified STRs on massively parallel sequencing platforms rather than electrophoresis can increase genotyping throughput (Scheible et al. 2014; Fordyce et al. 2015), although the scale of that analysis is still largely limited by the initial PCR amplification approach. Additionally, reduced-representation genomic datasets generated by restriction-site-associated DNA sequencing (RAD-seq) have been shown to yield useful, though limited in number, sets of discoverable STRs (Bonatelli et al. 2015), and genotyping-by-sequencing (GBS) methods have been effectively adapted for multiplexed recovery of known STR loci, again with a modest number of targets (Vartia et al. 2016). At the genome-wide scale, lobSTR (Gymrek et al. 2012) is a mature computational pipeline for high-throughput STR genotyping from massively parallel sequence data, and can be robustly deployed for personal genomic applications and large datasets of human genome sequence reads and those for other species for which a representative reference genome is also available. However, while lobSTR is a high-quality tool for accurate genotyping, it is not designed as a strategy for marker development or resequencing. MIPSTR (Carlson et al. 2015) is a recently developed pipeline for targeted STR resequencing using molecular inversion probes (MIP), with high reported marker fidelity and the ability to discriminate somatic variation from PCR slippage. However, this system also relies on a high-quality reference genome assembly, precluding use with the vast majority of species at present. Further, while the per-sample cost of MIP synthesis is modest if targeting very large numbers of individuals in a single species, the MIP cost advantage is gained by the effectively unlimited molarity of each probe following oligo synthesis. As such, it would typically be cost-prohibitive to target thousands of STR markers in a single study population using MIP. Thus while these and other purpose-built tools are robust and useful for many applications, there is not yet a targeted STR resequencing strategy with the ability to make cost-effective use of NGS data among diverse species for population genomics.

Some key goals of an STR resequencing strategy for diverse non-model species are i) no reliance on a reference genome, ii) cost-effective implementation even at single-project scales, iii) scalability to efficiently recover thousands of targets, iv) applicability to DNA from non-tissue sources such as fecal samples, and v) the simultaneous collection of STR and SNP genotype data to facilitate analyses of both datasets separately and/or in combination. We developed and tested a strategy to satisfy these criteria: Briefly, we developed a software pipeline, BaitSTR, to discover and locally assemble simple STR loci from unassembled NGS data without using a reference genome. We used the pipeline to design a biotinylated RNA bait library for in-solution capture (Gnirke et al. 2009), and to resequence and genotype 5,000 target STR loci in an endangered lemur. We tested the approach in both tissue (n=3) and fecal (n=1) samples. Further, we implemented a SNP discovery step in regions flanking the target STRs (in total, 1.2 Mb of targeted sequence), producing a large number of co-phased STR–SNP compound markers that could facilitate analyses focused across a wide range of spatiotemporal scales. Finally, we conducted a series of simulations to test the sensitivity of BaitSTR at various coverage levels and parameter permutations. In sum, the BaitSTR pipeline provides a cost-effective and scalable strategy for genomic STR discovery and targeted resequencing without the need for a reference genome or tissue DNA purity.

## Methods and Sample Materials

The BaitSTR software pipeline (Figure 1) consists of a modular set of programs to i) discover STRs through a rapid scan of short read shotgun sequence data, ii) collapse reads carrying STRs into a set of candidate loci, and iii) extend these candidates through a local assembly process to characterize the flanking sequence. The result comprises a large set of STR loci with regional sequence information from which DNA capture probes can then be designed based on user parameters. We tested the approach by first shotgun sequencing two diademed sifaka genomes each to medium coverage (∼12x), passing the shotgun data through the BaitSTR pipeline to select regions for targeting, and carrying out a capture experiment on three tissue and one fecal diademed sifaka samples. We also performed a set of simulation-based analyses to test the performance and parameter impacts of BaitSTR in the context of highly controlled datasets of known composition, using i) a random simulated “genome”, and ii) read data from the widely studied NA12878 CEPH genome.

**Figure 1:**
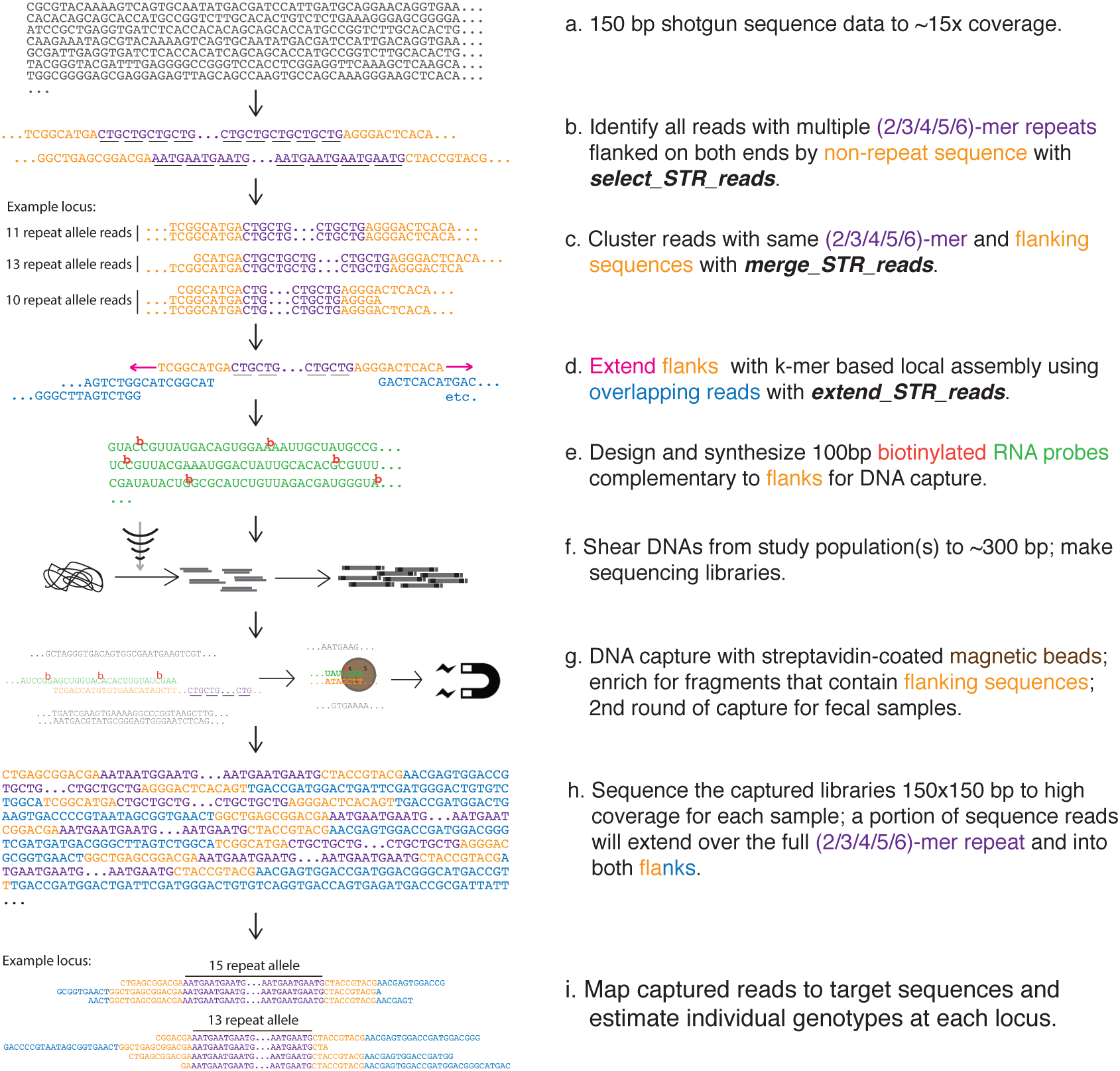
BaitSTR computational pipeline and massively parallel STR enrichment strategy.

### Marker Development

The BaitSTR software proceeds in three major steps; each executed using a separate module, for marker discovery and development (Figure 1):

#### Step 1. Identify input sequences harboring STR motifs

In a module named *select_STR_reads*, we implement a regular expression search to identify sequence reads harboring simple STRs. For example, we may identify reads containing all instances of 1nt to 6nt units repeated a minimum of five times in tandem; “CG CG CG CG CG” as a 2-mer STR, and “AAGCGT AAGCGT AAGCGT AAGCGT AAGCGT” as a 6-mer STR. Sequences may be optionally filtered to exclude homopolymer runs or may be constrained by the number of core repeat units and the length of the unique flanking sequences. We filter and retain these sequences, along with auxiliary information including the observed core repeat motif, the number of repeat units in the sequence, and the position of the STR in the sequence. In cases where more than one STR is found in the sequence, we keep the information about the STR that covers the most number of bases in the sequence. This step speeds up computations in subsequent steps, but also introduces a source of false-negatives by allowing discovery of only one marker per read. This module uses the ‘multiprocessing’ package in the Python programming language to implement data parallelism and leverage multiple cores common in modern CPUs. One of the inputs to this module is the length of the sequence flanks required around the STR on the discovery read, allowing for selection of STRs that are away from the ends of the sequences to ensure sufficient non-redundant flanking regions for downstream steps. However, it also has the adverse effect of increasing the false-negative rate by limiting discovery of STRs that effect less than L-2f bases of the sequence, where L is the length of the read, and f is the user-input flank requirement in base pairs. We also use the flanking sequence along with the STR core unit to flag candidate sequences that could support the same STR location as described in the next step.

#### Step 2. Merge reads supporting the same STR

In the module named *merge_STR_reads*, we collect all the sequences annotated in the previous step (*select_STR_reads*) that support the same observed core repeat unit or motif, and have flanking sequences that are similar as determined using alignment identity thresholds. As a sequence identified and retained from the previous step is read, the non-repeat flanks of the annotated STR are aligned to the flanking sequences of all the STRs that have the same repeat unit. We use a simple implementation of the Smith-Waterman algorithm (Smith and Waterman 1981) to perform the alignments of the flanks, and a sequence is added as evidence for an existing STR if the identity of the alignment and the block (a collection of reads supporting the same STR motif identified from previous reads) exceeds 90% and allows no more than two gaps. Consensus sequence for the segment flanking the STR is updated as more sequences are added as evidence for that STR. If the number of sequences in the block exceeds statistical estimates for the read coverage in a whole genome shotgun sequencing experiment via the Poisson approximation to the Binomial distribution (Lander and Waterman 1988), it is considered to be indicative of collapsed repeats or erroneous micro-assembly around the STR, and hence filtered out during this step. The user can also input a coverage threshold that overrides this default setting. Only blocks that show evidence of support for the presence of two different alleles, and hence polymorphic STRs, are output by default. However, a user-specified option allows retention of all the STR blocks, which may be desirable in cases where the discovery of STRs is performed in one individual and then genotyping is done in others.

#### Step 3. Extend STR contigs on both ends

We use the module *extend_STR_reads* to create local assemblies. This step extends the flanking regions to serve as probe targets and prevents inadvertently targeting (non-STR) repetitive elements or low-complexity regions. We extend the unique regions flanking each of the previously identified STRs using a *k*-mer based approach. Similar to BFCounter (Melsted and Pritchard 2011), we use a Bloom filter to probabilistically identify and store all non-singleton *k*-mers in a memory-efficient hash table implementation (https://github.com/google/sparsehash.git). This approach limits the inclusion of singleton *k*-mers arising from sequencing error while managing the intensive computational footprint of *k*-mer hashing. A second round of counting is done to remove *k*-mers that are only seen once (or seen below a user-specified threshold number of times) but are included as a result of the probabilistic nature of Bloom filters. We use the resulting list of non-singleton *k*-mers to extend both flanking sequences for an STR as follows: The leading *k*-mer is selected and possible extensions in the hash table are evaluated. If only one extension is found in the list of non-singleton *k*-mers, then that extension is applied and the process repeats. If two extensions are possible and the location represents a simple SNP, we can optionally tolerate the polymorphism by selecting the first of the alleles sorted lexicographically and extending the flank. If a more complex case is encountered, such as the edge of a repeat segment or a structural variant, then we terminate the extension. By default, the flanks are extended up to 1,024 bases on both sides of the STR, but the user can override this default by invoking a “--flanks” command line option. Extensions are also stopped if the next *k*-mer has already been used in extension of the same STR earlier during the extension process, potentially indicating a tandem repeat in the flank.

#### Diademed sifaka sequencing and STR genotyping

To test the BaitSTR pipeline, we first shotgun sequenced two diademed sifakas (male “Oberon” and female “Tatiana”—now-deceased captive individuals at the Duke Lemur Center who were wild-caught as adults in Madagascar) for marker development. DNA was isolated from liver tissue using Qiagen DNeasy Blood and Tissue kits, sheared to ∼325bp fragments using a Covaris S2 Ultrasonicator, and size-selected to the same target range on a 2% agarose gel. We targeted ∼325bp inserts for size selection to generate maximum non-overlapping data per insert while minimizing physical distance between forward and reverse reads. In post-capture libraries discussed below, this approach should increase the on-target density of sequence data by minimizing the proportion of insert regions captured but not sequenced, but insert size and sequencing configuration could be co-optimized in other ways. Illumina sequencing libraries were prepared after Meyer and Kircher (2010) with 12 cycles of indexing PCR using KAPA HiFi DNA polymerase. Both libraries were sequenced on one lane each of an Illumina HiSeq 2500 in Rapid Run mode with 300 cycle paired-end configuration (i.e., 150×150 bp reads), yielding an estimated 12x average depth of non-redundant coverage per individual (based on a genome size estimate of 3 Gb, i.e., similar to that of other lemurs (Perry et al. 2012b)). The sequencing run from Tatiana’s library produced reads with an elevated proportion of failed cycles, yielding data suitable for genotyping but possibly not for marker development. Marker development was therefore carried out using reads from Oberon only, invoking the option to output all STR blocks including both homozygous and heterozygous STR locations. The data from Oberon were filtered to exclude reads with any ambiguous base calls (Ns), leaving 260,598,672 reads that were carried through the STR discovery pipeline described above. *select_STR_reads* was configured to recover 2-6mer motifs with at least 5 tandem repeats and 27nt of flanking non-repeat sequence, yielding 4,014,698 STR-containing reads. 475,115 putative STR loci remained after read merging under default parameters using *merge_STR_reads*. Finally, 27-mers were used for block extension into each flank, SNPs were not tolerated during extension, and *k*-mer representation was capped to 30x using *extend_STR_reads*.

#### Probe design and target capture

Shotgun reads from both sifaka individuals were then re-mapped to this set of blocks using parameters identical to the BWA-backtrack strategy in the *BaitSTR_type.pl* script, and genotyping was carried out at loci with at least 4 independent, non-redundant reads with 30nt flanking alignments and an integer value for repeat number (i.e., no partial repeats). 63,596 STR locus genotypes were called in Oberon and 57,745 were called in Tatiana, with 55,531 loci in common. We discarded loci with more than two alleles present in either individual, suggesting PCR stutter or somatic variation (n=4,915 discarded), and we increased the minimum repeat number for 2mers to 7 to further pare down the target set, eliminating 35,131 further loci. We then selected a custom set of loci with at least 2 total alleles observed across 4 chromosomes, totaling 3,882 sites (1,041 2mers; 1,803 3mers; 908 4mers; 90 5mers; 40 6mers). To reach our target of 5000 sites for capture, we selected an additional 1,637 3mers (n=1,074) and 4mers (n=563) with only one allele observed, but with a minimum 6 repeats. We designed four individual 100mer probes per target: two on each flank, with one probe per flank immediately abutting the STR, and another probe per flank 20nt away from the STR, overlapping the first probe by 80nt. All four probe sequences per targeted locus were required to pass the probe design QC pipeline implemented by MYcroarray to retain a locus. 3,505 of the 3,882 multi-allele sites passed probe design QC, so 1,495 of the 1,637 single-allele sites were included at random (the 5,000 chosen target sequences are provided as supplemental dataset S1). We used lastz (Harris 2007) to identify the orthologous regions of 5,000 target regions in the hg38 human reference genome assembly (where possible given sufficient sequence conservation). We directly aligned the complete target blocks to the hg38 assembly using default lastz parameters. We then filtered alignment matches to a minimum 50% coverage of the target and 70% sequence identity, and retained targets with only one valid match across the human genome. This strategy recovered the orthologous locations of 4,249 (85%) of the sifaka STR loci. Of the mapped loci, 2,267 overlapped annotated genic regions (including introns) and 1,982 were entirely intergenic, and excluding the Y chromosome (our preferential selection of variable markers would have limited the inclusion of such markers; the initial discovery step could include more male individuals in alternative designs for future studies) the markers are distributed across the genome (Figure 2).

**Figure 2:**
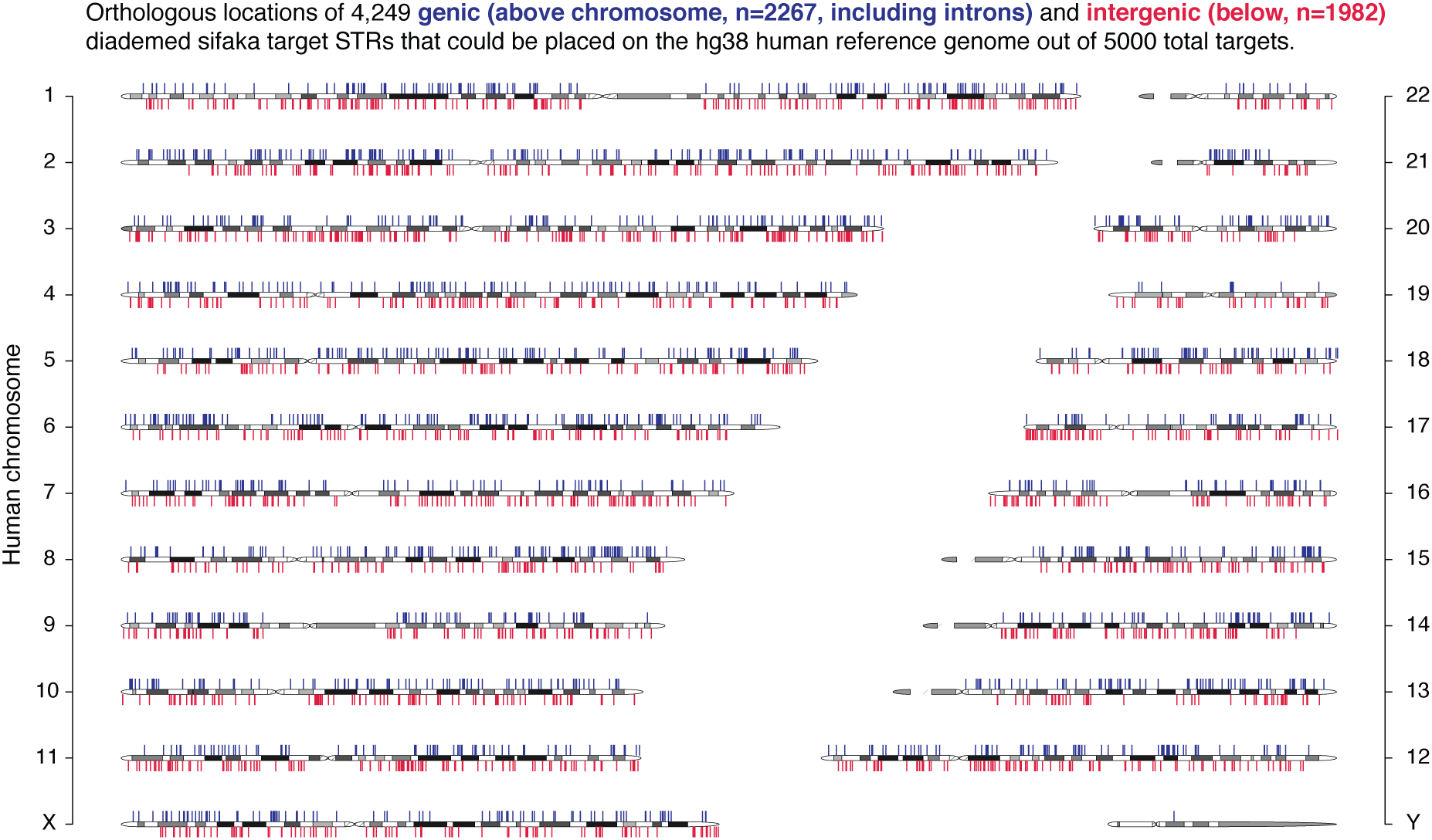
Orthologous locations of diademed sifaka STR targets on the human reference genome. A) Genome-wide distribution of genic (blue, n=2,267) and intergenic (red, n=1,982) diademed sifaka STR loci that could be mapped the human genome.

The probe biotinylation process for target immobilization on streptavidin beads (Gnirke et al. 2009) is implemented through the integration of biotin-UTP at genomic thymine sites in the probe sequences, so a moderate probe uracil content is required for effective target recovery. Probe sequences with fewer than 11 Us were discarded, and 301 probes with low U content (11–14 Us) were given U-tails of 1–5 residues to ensure effective biotinylation. The resulting 20,000 probes were obtained from MYcroarray in a MYbaits-1 kit (probe design lot 140715-31; probe sequences provided as supplemental dataset S2).

We prepared additional sequencing libraries from two further samples: blood and fecal DNA from “Romeo”, the male offspring of Tatiana and possibly Oberon, but with uncertain paternity. Romeo was wild-caught as a juvenile along with his mother Tatiana; Oberon had been captured previously from a different group in the same population. Romeo (now deceased) survived Tatiana and Oberon at the Duke Lemur Center, making possible the collection of both blood and feces from this individual. The fecal samples used in this study were stored in 10 mL RNALater upon collection and stored at ambient temperature until received in the laboratory several days following collection, after which these samples were stored at at −80° C. Blood DNA for Romeo was extracted from 50µL whole blood using a Qiagen DNEasy Blood and Tissue kit following the manufacturer’s protocol. DNA was extracted from 200mg of fecal material per extraction. Following previous fecal DNA genomics capture research (Perry et al. 2010), we combined separate fecal extractions to reach a total of 8µg of total DNA (i.e., including endogenous DNA from the individual, along with presumably a much larger quantity of exogenous DNA, for example from gut bacteria (Perry et al. 2010)) per individual, then concentrated the DNA samples using a SpeedVac to yield three separate aliquots of 130µL for shearing for each individual. Fecal aliquots were then sheared on a Covaris M2 according to the manufacturer’s protocol for 350bp sheared products, with the time reduced from 65 sec to 60 sec to correct for observed slight overshearing of fecal DNA. 2µg of blood DNA was sheared to 350bp in a 130µL aliquot according the recommended Covaris M2 protocol. Following the fecal capture protocol described by (Perry et al. 2010), sheared fecal DNA was concentrated to 2-3 aliquots of 25 µL per sample and size-selected on a 2% low-melt agarose gel to ∼350bp. Gel-extracted DNA was further purified using SPRI beads prior to enzymatic steps of library preparation. Barcoded sequencing libraries were prepared from these samples using TruSeq Nano kits (Illumina) following the manufacturer’s protocol for a 350bp insert size using 8 PCR cycles. Two duplicate libraries were made from the combined isolated fecal DNA, using different barcodes.

All five libraries—the two original shotgun-sequenced samples from Tatiana and Oberon, one new tissue library from Romeo, and two duplicate libraries of fecal DNA from Romeo—were enriched for the STR targets using the MYbaits protocol version 2.3 (MYcroarray). For blood and tissue libraries, 500ng was used for hybridization. For fecal samples, the input DNA was increased up to 1.3µg, and all protocol reagents besides the capture master mix were doubled, according to previous modifications (Perry et al. 2010). Hybridization was carried out for 24 hours at 65° C, and the number of post-hybridization kit washes was increased from three to six based on previous procedures (Perry et al. 2010). The resulting templates were each re-amplified in two separate reactions for 14 cycles according to the MYbaits PCR protocol using library primers IS5 and IS6 (Meyer and Kircher 2010) and KAPA HiFi DNA polymerase. Enriched libraries were re-combined immediately after amplification and purified with SPRI beads. Using one of the two enriched duplicate libraries of fecal DNA, we carried out a second capture using 300ng of the enriched library as input. Reagents were doubled as above, and re-amplification PCR was carried out for 11 cycles. We sequenced the enriched libraries in multiplex pools on partial lanes of either HiSeq 2500 or NextSeq instruments with 300 cycle paired-end reads, recovering between 17.4 million and 36.5 million read pairs per library (sequencing details provided in Table 1).

Sequence reads were aligned to the targets using *BaitSTR_type.pl*, a companion script to BaitSTR (https://github.com/lkistler/BaitSTR_type). *BaitSTR_type.pl* can use the complete lobSTR pipeline (Gymrek et al. 2012), lobSTR with a BWA-mem alignment (Li 2013), or a BWA-backtrack approach (Li and Durbin 2009) for read mapping and STR genotype calling, and includes all pre-processing and reference indexing steps to filter and prepare the output blocks from BaitSTR. The BWA-backtrack pipeline was developed to accommodate divergent samples with a previous version of lobSTR that was not yet compatible with external aligners, given lobSTR’s limitation with divergent read mapping. It was used to design the target markers and is included with *BaitSTR_type.pl*, but should now be considered deprecated. We used the BWA-mem/lobSTR pipeline to align the captured reads to the complete extended blocks of the target 5000 loci (500–2276nt each block, total 5.68Mb) and call STR genotypes. We used default parameters with the following changes: We modified genotype calling parameters in the *allelotype* module (Gymrek et al. 2012) as recommended in lobSTR documentation for handling BWA-mem alignments (--realign --filter-clipped --min-read-end-match 10 --filter-mapq0 --max-repeats-in-ends 3). We carried out PCR duplicate removal using the SAMtools 0.1.19 *rmdup* function (Li et al. 2009), separately collapsing the mated and unmated reads (default behavior in *BaitSTR_type.pl* with bwa-MEM alignment), and disabled duplicate removal in *allelotype* (--normdup). SAMtools *rmdup* default behavior fails to collapse unmated paired reads. However, in the case of mapping to small targets, a high proportion of read pairs contribute only one of the two reads to the alignment, potentially leaving a large set of PCR duplicates in the alignment, whereas treating paired reads as single reads artificially collapses inserts with differing outer coordinates. In the shotgun data from Oberon, unmated reads constitute 58% of the aligned data. As such, these unmated reads are separated and treated as single reads, then recombined with the mated pairs and re-sorted.

## Results

### STR enrichment

Briefly, we shotgun sequenced two diademed sifaka individuals and designed an STR bait library using the BaitSTR pipeline. We then tested STR target capture by RNA hybridization using three tissue DNA samples and one fecal DNA sample. The three tissue libraries were highly enriched for the target loci compared to shotgun datasets, with a large proportion of captured regions yielding suitable non-redundant sequence read coverage for genotype calling (Figure 3). Given the composition of our target STR loci, there are 498,606 genomic positions from which 150nt reads requiring 15nt non-repeat flanks could be initiated and yield a genotype call, predicting that 0.000156 of shotgun reads should yield target genotype calls in a 3.2Gb genome. In the shotgun data from Oberon and Tatiana, we observed a slightly lower proportion of callable reads than this genomic expectation (.000128 and .000118, respectively), which is likely due to PCR redundancy and read filtration. In the capture data from these two individuals, we observed .0187 and .0127 STR-callable reads—reads completely spanning the STR according to the parameters described above for *allelotype* (Gymrek et al. 2012)—constituting empirical enrichments of 145x and 109x, respectively. In total, the three tissue samples yielded 4710 (94.2%; Romeo), 4939 (98.7%; Oberon), and 4953 (99%; Tatiana) of the 5000 target loci callable at minimum 10x non-redundant coverage—or with at least ten independent sequence reads spanning the STR for genotype calling. Assuming a 100-fold enrichment factor, one Illumina HiSeq 3000/4000 lane producing 87.5Gb of read data would thus be predicted to yield an average of 37x coverage of the target regions for each of 96 samples sequenced in parallel.

**Figure 3:**
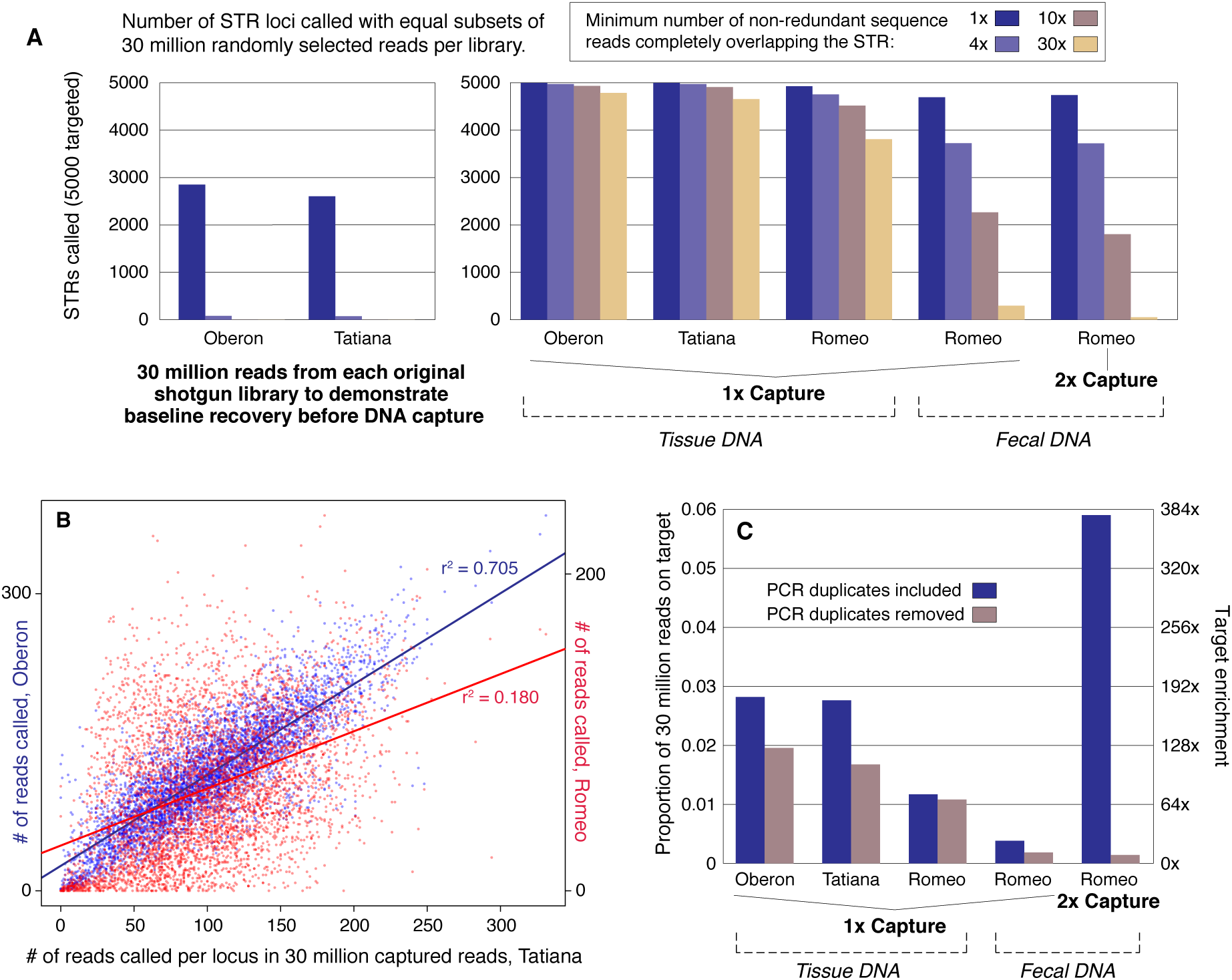
Target Enrichment Results. A) Number of target STR loci recovered in shotgun (n=2) and captured (n=5) libraries. Read data were randomly downsampled using SAMtools (29) after read mapping and before genotype calling to normalize all libraries to 30 million input reads for cross-comparability. Actual reads generated per captured sample ranged from 34.8 million to 73 million (Table 1). B) Per-site coverage is highly correlated among samples, illustrating non-random variation in marker enrichment. Marker coverage in the 30 million read subsample is compared between Tatiana Oberon capture data (left axis, blue), and Tatiana and Romeo’s tissue library (right axis, red). C) Enrichment of reads carrying target STRs in subsamples of 30 million reads. Left axis shows the proportion of callable reads, right axis shows the estimated enrichment level given the genomic expectation of 0.000156 of reads on target with no enrichment. For the fecal libraries, enrichment values are given without any correction for the high proportion of exogenous DNA, whereas previous estimates of endogenous fecal DNA content (Perry et al. 2010) suggest actual enrichment similar to the tissue samples.

We randomly subsampled all read alignments to approximately 30 million input reads using the SAMtools *view* function with the “-s” option (Li et al. 2009) to compare results between samples (Figure 3). Coverage was found to be highly correlated between samples, with Oberon and Tatiana’s capture data showing a strong linear relationship in per-locus coverage (Pearson’s product moment correlation: r^2^ = 0.70, p < 2 × 10^−16^; Figure 3B). Romeo’s per-locus coverage also correlates significantly with Oberon (r^2^ = 0.31, p < 2 × 10^−16^) and Tatiana (r^2^ = 0.18, p < 2 × 10^−16^), but lower correlation coefficients likely reflect different biases introduced during the two different library preparation strategies used (Seguin-Orlando et al. 2013). Moreover, in the subsampled data, the three tissue capture samples yielded 4933 (Oberon), 4909 (Tatiana), and 4516 (Romeo) marker genotypes at 10x coverage or greater. Thus, at random, ∼4374 10x genotyped markers would be expected to be common among the three, but instead we observed 4497 10x genotyped markers in common. This result further demonstrates that marker dropout occurs systematically rather than at random, likely as a function of hybridization efficiency, which is advantageous in terms of retaining overlapping genotype calls between samples in larger test populations. That said, in general the overall bias in terms of marker coverage introduced by the capture process was subtle, so that this systematic dropout does not lead to major unevenness in marker coverage across samples.

Our method also successfully enriched Romeo’s fecal DNA test libraries, although the poorer DNA quality and the high proportion of exogenous DNA in these samples did affect the genotyping process as expected (Perry et al. 2010). Following one round of capture, 4815 markers were called with at least one read in the entire dataset before subsampling to 30 million, and 3858 were present at the 5x level or greater. We recovered .00113 of reads yielding genotype calls, representing 7.27x “effective” enrichment—that is, given genomic expectations of .000156 in a completely endogenous template. However, fecal samples are dominated by exogenous DNA, with major contributions from gut microbiota and other sources (Perry et al. 2010). Assuming a conservatively high estimate of 5% endogenous DNA content in feces (Perry et al. 2010), our results would constitute an actual 145x enrichment of the STR targets, which would be on par with enrichment levels observed in the tissue samples. Previously, a serial capture approach was used to further boost the on-target proportion of reads from fecal libraries (Perry et al. 2010), and so we tested a second capture with one of the fecal libraries. In our case, serial capture increased dramatically the proportion of on-target reads prior to PCR duplicate removal from 0.00381 to .0628, a 16-fold increase, but the added PCR cycles ultimately lowered library complexity to a problematic degree. After removal of duplicate reads, the on-target proportion of reads declined slightly to .000738. Thus in this case, the serial capture approach yielded fewer genotypes than a single capture due to a loss in library complexity.

### Familial genotyping

As a test of STR fidelity, we used KING (Manichaikul et al. 2010)—which computes a kinship coefficient reflecting the degree of relatedness on the basis of genomic data—to test for relationships among the putative familial trio of Tatiana, Oberon, and Romeo using the genotype. The kinship coefficient statistic calculated by KING in the ranges of >0.354, between 0.177 and 0.354, between 0.0884 and 0.177, and between 0.0442 and 0.0884 correspond to duplicate/monozygotic twin, 1st-degree, 2nd-degree, and 3rd-degree relationships, respectively (Manichaikul et al. 2010). Tatiana is the known mother of Romeo, while Oberon’s paternity was suspected but uncertain. The KING analysis reported a kinship coefficient of 0.2974 between Romeo and Tatiana, confirming a first-degree relationship between the two consistent with either parent-offspring or full sibling relatedness (Manichaikul et al. 2010). However, results from KING were inconsistent with 1^st^ or 2^nd^ degree familial relationships between Oberon and Romeo (kinship coefficient = 0.0519). Given the robust result supporting Tatiana’s known parenthood of Romeo, we can conclude that Oberon is not the father or a close relative of Romeo. Our method’s high genomic resolution holds substantial potential for determining relatedness within groups and populations for which relationship structures cannot be fully understood from behavioral observation, as is often the case for *Propithecus* species, for which there is both variation in the number of adults of each sex within social groups and one previous study identified a high rate of extra-group paternity (Richard 1985).

### STR-linked SNPs

The library capture approach generates a large amount of resequencing data beyond the direct genotype information at STR loci, and therefore also functions as a reduced-representation genomic dataset with a substantial number of incidentally captured single nucleotide polymorphisms (SNPs) across samples. These data could be utilized for a wide range of analyses, either as an independent data class or in conjunction with STR data. In fact, compound markers consisting of an STR and one or more tightly linked SNPs are potentially useful for high-resolution population genomic applications (Agrafioti and Stumpf 2007), and the incidental capture of SNP panels during STR targeting yields numerous of these linked markers (Figure 4). We used the SAMtools *mpileup* function (Li et al. 2009) and VarScan (Koboldt et al. 2009) to screen the alignment files produced by *BaitSTR_type.pl* for SNPs with at least 4x coverage per sample. We excluded SNPs within the targeted repeat region itself, and we called SNPs only where they could be linked directly to a called STR allele, either on the same read or on a mated read, facilitating secure phasing. This approach identified 13,955 SNPs in total, including 13,005 genotyped (with minimum 4x coverage) in all three tissue samples. Increasing the coverage threshold to a 20x minimum, we retain 10,715 SNPs that were genotyped in common among the three tissue samples. By relaxing the requirement for co-phasing with STRs through co-occurrence on an insert, we recover a total of 21,909 SNPs, 18,520 of which are present in all three tissue libraries at 20x coverage or greater.

**Figure 4.**
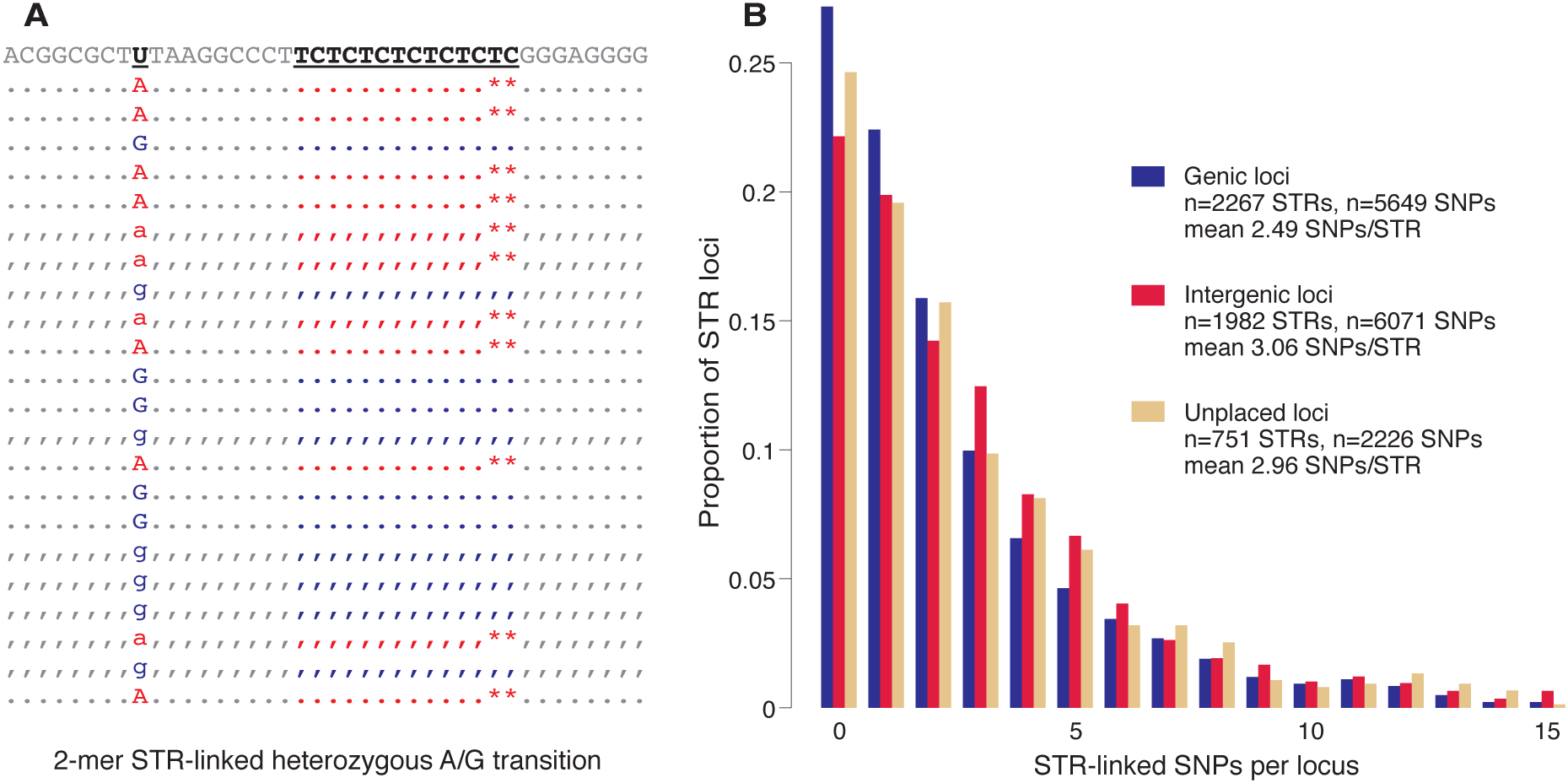
SNP-STR compound markers. A) Simulated example of a phased STR-linked SNP locus, where the ‘A’ SNP allele associates with six repeats of the TC motif and the ‘G’ allele associates with seven. B) Proportions of genic, non-genic, and unplaced STR loci (based on mapping analysis to the human reference genome; see Figure 1) with number of SNPs detected at ≥4x coverage on associated inserts from three lemur tissue sample libraries (Oberon, Tatiana, and Romeo). 42 STR loci associated with >15 SNPs (n=8 genic, n=25 non-genic, n=9 unplaced) are not shown.

Diademed sifakas have the highest level of mitochondrial genome sequence diversity among a sample of eight extant and two extinct lemur species for which population mitogenomic data are available (Kistler et al. 2015). However, the STR-linked SNP data that we generated for diademed sifakas in this study could not be used for comparison to nuclear genome estimates of nucleotide diversity to those available for other lemurs (Perry et al. 2012a), because our STR marker selection (based on Oberon’s shotgun sequencing data) excluded loci with evidence of flanking region SNPs, thereby also artificially reducing the observed number of heterozygous sites in other individuals with shared population history to Oberon (i.e., Tatiana and Romeo). While we have subsequently modified BaitSTR to allow users to select STRs irrespective of the presence of flanking region SNPs in the individual(s) on whose shotgun genome sequence data those steps were performed, this experience does illustrate the importance of considering the effects of various study design options. For example, if at the marker development step the user chooses to preferentially select variable STR loci, then the individuals whose shotgun genome sequence data were used for that step should be excluded from population genetic diversity analyses of any type of marker in to avoid inflation of diversity statistics.

### PCR Stutter

PCR “stutter, ” the physical slippage of DNA polymerase on the template strand resulting in loss of repeat fidelity (Schlötterer and Tautz 1992), is a known obstacle for traditional PCR-based STR analysis (e.g. Hoban et al. 2013). We found that PCR stutter was ubiquitous in our captured libraries, due to both the initial library amplification and the subsequent reamplification step required after target capture to reach molarity sufficient for cluster generation. In our test dataset, we found that PCR stutter can influence genotype calls, but the effects are manageable. We tested for the effect of PCR stutter by comparing i) Oberon’s shotgun and capture data, expected to match, ii) Tatiana’s shotgun and capture data, expected to match, iii) Romeo’s tissue and fecal capture data, expected to match, and iv) Tatiana and Romeo’s capture data, expected to share at least one allele in common. We categorized markers into three groups for each pairwise comparison between libraries: i) full matches at both alleles, ii) ‘half matches’, where one allele matches and the other does not match (this category includes markers where one library appears homozygous and the other appears heterozygous with one ‘allele’ matching the homozygote), and iii) mismatches, where neither ‘allele’ matches between libraries, with libraries appearing as any configuration of homozygous and heterozygous. Further, PCR stutter usually skips or adds only one repeat period at a time, yielding a decay profile peaking at the true biological allele (Gymrek et al. 2012). Therefore we further divided half-matches into ‘stutter half-matches, ’ where the mismatching ‘allele’ differed by only a single repeat period, and ‘non-stutter half-matches’ representing any other configuration. Finally, we note that any half-matches with one ‘homozygous’ call might represent failure to observe a true heterozygote due to dips in coverage or recovery biases.

We first compared mother Tatiana and her offspring Romeo at 10x sites in common, where we expect either complete or half matches at all alleles. With a minimum of 10x non-redundant read coverage we expect both alleles at heterozygous sites to be represented >99.8% of the time based on binomial sampling probability, thus limiting false-positive PCR stutter identifications and false-negative matches from allelic dropout. On the basis of Romeo’s blood library and Tatiana’s capture dataset, 99.5% of sites satisfied this requirement of full or half matches (4676 out of 4701 sites present in both libraries at 10x). Similarly 99.7% of sites (2852 out of 2862) satisfied the requirement with Romeo’s lower coverage fecal DNA library.

We also compared the shotgun and capture libraries for consistency within a single sample for Oberon and Tatiana. These comparisons necessarily included fewer markers because of the lower overall coverage of the shotgun datasets. Nonetheless, out of 1338 markers compared between Oberon’s shotgun and capture data at a minimum 10x depth of coverage, 1233 (92.2%) were full matches at both alleles and none were full mismatches. Among the 105 half-matches, 97 (92.4% of half-matches, 7.2% of markers compared) are classified as ‘stutter half-matches’ as described above. The remaining 8 loci are ‘non-stutter half-matches.’ Representing only 0.6% of compared markers, these observations could reflect the small expected proportion of heterozygous sites for which both reads were no represented even at 10x coverage. A comparison of Tatiana’s 10x shotgun and capture genotypes (1413) yielded very similar results, comprising 91.3% matches, 8.4% stutter half-matches, 0.4% non-stutter half-matches, and a no full mismatches. Finally, when comparing 2860 markers called in Romeo’s tissue and fecal libraries, we observed 89.8% perfect matches, 8.6% stutter half-matches, 1.6% non-stutter half-matches, and no complete mismatches. A slightly elevated stutter rate is expected in this comparison given that both libraries were captured and re-amplified, unlike Oberon and Tatiana’s shotgun datasets.

## Simulations

In order to assess the limits of STR detection and assembly under variable levels of data coverage and *k*-mer size parameters, we performed several simulations to test the sensitivity and practical limitations of the BaitSTR pipeline.

### STR discovery in a simulated random genome

We simulated a 2 Mbp long chromosome of entirely random DNA to understand the constraints imposed by sequence coverage and *k*-mer length for automated STR discovery and extension using BaitSTR. We added bi-allelic STRs after every 2000 bp stretch of random sequence in this synthetic genome, and then ran our pipeline to investigate the recovery rate of the STRs. In the absence of repeat sequences, the rate of recovery of such STRs should be high, and only depend *k*-mer length and genomic coverage as follows:

- *k*-mer length. Sequences that harbor STRs with non-repeat flanking sequence at least equal to the *k*-mer length are required for STR discovery in *select_STR_reads*, so that the candidate region of reads for STR discovery decreases with larger *k*-mers given a constant read length. Additionally, *k*-mer length is an important factor in later stages, as the flanking regions are used to collapse candidate STR loci in *merge_STR_reads*, and *extend_STR_reads* uses a *k*-mer based extension process sensitive to variable *k*-mer lengths.
- Average coverage. This variable determines the number of reads that support a STR allele. By default, we require at least 3 reads to support a polymorphic STR location, before it becomes a candidate for subsequent extension.

We performed two sets of simulations with this dataset. First, we focused on understanding the limitations of our approach and the effects of the various design decisions that were made during the discovery step in *select_STR_reads.* To accomplish this goal, we ran the pipeline but did not impose any restrictions on the minimum length of the extended contigs prior to calculation of performance metrics. For the second simulation, we required that the extended contigs satisfy minimum total length of 500nt, and minimum non-repeat flanks of 200nt (similar to the requirements imposed for our analyses with real datasets) before calculation of performance metrics. In each of these simulations, Illumina short-read sequences were simulated using pIRs (Hu et al. 2012). The average coverage of the sequences was varied between 5-fold to 39-fold, and BaitSTR was run using *k*-mer lengths between 9 to 31 bps for each of those coverage values. In each of those runs we required a flanking sequence around the STR to be at least equal to the *k*-mer length used in that run. The extended contigs from *extend_STR_reads* were then aligned back to the reference genome using BLAT (Kent 2002). The alignments were processed to calculate the mapping locations of identified STRs on the reference sequence using an in-house custom script, and the true positives and false-positives were subsequently calculated using BEDtools (Quinlan and Hall 2010). False-positives are defined as i) chimeric contigs where extension resulted in incorrect local assembly, ii) contigs incorrectly aligned back to the reference, iii) collapsed repeats that could masquerade as polymorphic segments, or iv) STRs detected by the pipeline that were not explicitly introduced during genome simulation. Importantly, the fourth type are legitimately present at random in a simulated genome, and so are not strictly false-positives concerning pipeline specificity. For instance, 12 out of 4^10^ (1,048,576) possible 10nt DNA sequences are 2-mer STRs repeated 5 times, predicting ∼23 instances of random 10nt 2-mer STRs (95% CI: 14 – 33) in a 2Mbp random sequence. In a biological sequence, these would be correctly identified as true STRs rather than false positives.

As expected, our ability to recover STRs in the random genome increases with coverage (Figure 5). Higher coverage allows for an increased probability that STR alleles at a location fall within the acceptable region on the reads and are supported by sufficient number of reads to become candidates for extension. Increasing *k*-mer length initially increases, and then decreases our ability to find the STRs, similar to the expectation in genome assembly using De-Bruijn graphs (Chikhi and Medvedev 2014). We also find that the false positive rate of our pipeline is low, and falls to zero at *k*-mer lengths higher than 9. That is, given sufficient *k*-mer length, all contigs are correctly assembled and placed. It is important to note that the reference in this case is composed of random sequences and is only 2 Mbp long, so significantly greater *k*-mer length would likely be necessary in the case of mammalian genome size and complexity. Roughly 10% of STRs are not recovered in these simulations. As described above, we select and store information for the STR that impacts the most number of bases in a sequence using *select_STR_reads*, and in cases with multiple STRs are detected in the same input sequence, this can lead to a situation where one of the alleles for a motif is always missed. Such a situation is more likely for 2mers with small numbers of repeats even in a random reference genome, compared to larger STRs or STRs with longer motifs. An analysis of the false-negatives reveals that they are indeed enriched for 2mers with smaller number of repeats. This result informs us of one of the current shortcomings of our computational approach. A polymorphic STR is missed if one of the alleles is not discovered; i.e., in those cases it is found to be homozygous for an allele, which can happen in situations where there is another STR within a distance less than a read length away from it. One improvement that could be explored would be to store information about all non-overlapping STRs in the first step, followed by subsequent selection of the motif that results in a polymorphic marker in the second step.

**Figure 5.**
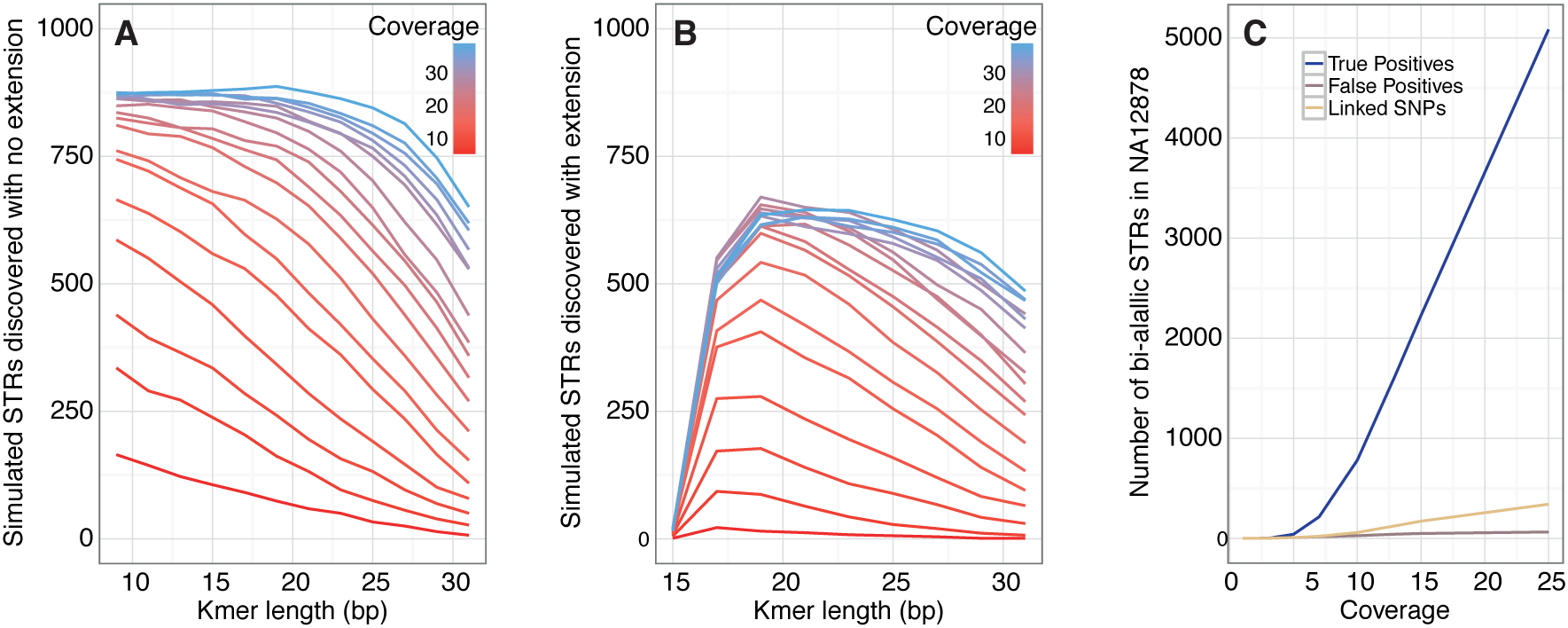
Simulated performance of STR discovery using BaitSTR. A) At variable *k*-mer lengths and coverage levels, the number of discoverable simulated bi-allelic STRs in a random synthetic genome that could be discovered with no requirement of local extension. 1000 total markers were present. B) Using the same simulated STRs in a synthetic genome, this simulation required successful block extension to 200nt non-repeat flanks and a total contig length of 500nt. C) At variable coverage levels, the number of heterozygous STRs discovered in the NA12878 genome data, along with a low frequency of false positives and the number of heterozygous SNPs recovered from extended blocks.

A key requirement of marker development is to identify STR locations with suitable flanking non-repeat sequence for targeting with RNA probes. The above simulation does not impose any extension requirements on the extended contigs, but marker development imposes a restriction that only bi-allelic STR locations satisfying criteria on length and uniqueness of the flanking region be selected for downstream processing. In order to assess these limitations, we also ran the above simulation at varying coverage requiring that the extended contigs satisfy minimum overall contig length of 500nt and minimum non-repeat flanks of 200nt, reflecting the requirements imposed for our lemur marker development. As expected, the false-positive rate remains 0, and the number of recovered STRs is constrained as above by *k*-mer length and depth of coverage (Figure 5). This requirement inevitably excludes some true STR loci that co-occur with repeat regions or difficult extension targets (low-complexity or highly heterozygous regions), but omitting these targets is important for in-solution capture to preempt saturation of the probes by repetitive reads. That is, only high-quality, securely non-repetitive markers should be targeted.

### STR discovery in human genome sequence read data

To analyze the detection performance of BaitSTR in a non-random genomic landscape, we used previously published shotgun Illumina sequencing data from the widely studied human NA12878 CEPH individual. We downloaded the read data alignment to the human reference assembly hg19 from ftp://ftp.sra.ebi.ac.uk/vol1/ERA172/ERA172924/bam/NA12878_S1.bam, comprising 791,385,507 sequence pairs resulting in ∼50-fold coverage of the human genome. We used SAMtools (Li et al. 2009) to identify and genotype putative STRs using the variant calling workflow for WGS described at http://www.htslib.org/workflow, followed by filtering to only keep putative STR variants. We converted the downloaded BAM alignments to raw FASTQ sequences using bam2fastx script supplied with TopHat (Trapnell et al. 2009), and then used lobSTR (Gymrek et al. 2012) to identify, align, and genotype all putative heterozygous STRs in the individual using the lobSTR index for hg19. We were able to genotype 1,518,346 of the putative STR locations using lobSTR, out of which 692,499 were non-homopolymers repeated a minimum of five times in tandem, 97,064 were called as heterozygous in NA12878, and 61,343 were exact repeats in at least one of the two genotyped alleles in NA12878. Importantly, BaitSTR targets only simple STRs whereas lobSTR detects complex motif combinations that include non-exact repeats, so that the all members of the lobSTR set may not be detectable using BaitSTR. We created a combined file in BED format that was used to represent the set of STRs found using SAMtools or lobSTR, and we treated them as the “true” set of calls for the following analyses.

We measured the detection capabilities of BaitSTR with the NA12878 genome resequencing dataset by randomly resampling read datasets with sequencing coverage ranging from 1-fold to 25-fold (Figure 5). We required a flank of 29 bases during STR discovery (*select_STR_reads*), and used an extension *k*-mer length of 27 (*extend_STR_reads*). As above in the second simulation with the random reference genome, we required extended contigs to contain 200nt non-repeat flanks with a minimum 500nt overall length. The assembled contigs were aligned back to the human reference using BLAT (Kent 2002), and only the alignment with the highest score was included to identify the putative STR locations on the reference. This set of aligned contigs was then compared to the “true” set of calls using BEDtools (Quinlan and Hall 2010). As expected, the STR recovery rate increased with coverage, as more STRs gain sufficient support and fall within the valid window for annotation and extension. At 25-fold coverage of the reference, we were able to recover 5,149 polymorphic markers, 5,085 of which overlapped with the “true” set of STR calls identified using a reference sequence by lobSTR and SAMtools— those markers that were heterozygous in NA12878 and could be extended on both sides of the STR with at least 200nt on either side with a total contig length of 500nt or greater. Further analysis showed that the calls at lower coverage are enriched for false-positives, and the fraction of false-positives decreases with increasing coverage. The false positive rate drops to less than 2% at 15x coverage. False-positives in this case are defined to be STR locations identified as polymorphic using BaitSTR, but which are not called as polymorphic by lobSTR or SAMtools. Although BaitSTR detected only 8.3% of potentially detectable polymorphic STR loci, our pipeline is designed to be conservative with respect to the risk of non-specific hybridization. Specifically, by implementing strict constraints on coverage, contig and flank size, and flank polymorphism, we limit the risk of inadvertently targeting non-STR repeat regions and low-complexity sequence for enrichment. Otherwise, the sequencing library could be swamped by repetitive element reads, which would decrease dramatically the effective enrichment of single-copy targets. As such, our pipeline errs on the side of avoiding this outcome, although user-defined inputs can be used to modify these variables with respect to target composition and specificity.

We used the NA12878 data at various coverage levels to measure the capability of BaitSTR to identify and genotype compound markers that include both an STR locus along with one or more tightly linked single nucleotide variants (SNPs). We only considered SNPs that could be linked to an STR allele on the same read to facilitate phasing. The raw sequences were aligned to the assembled contigs using BWA, and the SNPs were called using SAMtools (Li et al. 2009) with analysis only of those sequences that also covered the polymorphic STR marker (i.e., such that the SNP allele is empirically phased with the STR allele; See Figure 4A). We removed SNPs with a Phred-scaled SNP quality score less than 10 as calculated by SAMtools, as well as cases where we did not find at least 2 reads supporting each allele. We found that the number of STR-linked SNPs also increase with increasing coverage, and we find such tightly-linked SNPs associated with 6.66% of the STRs in this dataset at coverage of 25x.

## Discussion

Our approach facilitates the robust identification of STR loci from short read data for species without a reference genome, enrichment for thousands of those STR targets from both high-quality and non-invasive samples using hybridization capture, and genotyping of both the targeted STR markers and nearby SNPs. From the data we generated for three diademed sifaka individuals with, we could correctly recover the parent-offspring relationship between two of the individuals (mother Tatiana and child Romeo), and reject a parent-offspring relationship with the third individual and the child (Oberon is not the father of Romeo).

In future implementations of BaitSTR, we recommend steps to mitigate PCR stutter, with care taken to limit the number of PCR cycles to an absolute minimum. For example, we would recommend the use of PCR-free library preparation kits, which are commercially available from several manufacturers. A round of PCR following target capture is currently indispensible to reach sufficient sequencing molarity, so we additionally recommend maximizing input DNA to the probe hybridization step and using qPCR quantification approaches to optimize the number of post-capture PCR cycles performed (e.g., after ref. (Meyer and Kircher 2010)). Library reamplification enzymes and PCR conditions could also be optimized to improve STR fidelity (Seo et al. 2014).

Our strategy for massively parallel STR discovery and resequencing is scaleable, cost-efficient, and appropriate for non-model species and non-invasive samples. For these reasons, it is suited especially to ecological genomics and conservation contexts where, for example, fecal samples are a particularly useful source of DNA from endangered and otherwise difficult-to-study species. This approach could be applied to studies of population genetic structure, the relationship between relatedness on behavior, and mate selection. In populations suffering from habitat loss and fragmentation, genomic-scale STR datasets could be analyzed to rapidly assess changes in population genetic structure and outbreeding due to discontinuity of habitat, and as necessary assist in the active management of small and/or subdivided populations.

## Acknowledgments

We thank the Duke Lemur Center for providing the samples. This is Duke Lemur Center publication no. XXXX. This work was funded in part by a grant from the National Science Foundation (BCS-1554834, to G.H.P.). Any opinions, findings, and conclusions or recommendations expressed in this material are those of the authors and do not necessarily reflect the views of the National Science Foundation. Instrumentation was funded by the National Science Foundation through Grant OCI–0821527. L.K. was funded in part by the Natural Environment Research Council (NE/L012030/1, to L.K.). We thank Craig Praul and Candace Price from the Pennsylvania State University Huck Institutes DNA Core Laboratory and Xinmin Li at the UCLA Clinical Microarray Core for assistance with sequence data collection. We thank Melissa Gymrek for helpful discussion about lobSTR.

## References

Agrafioti I, Stumpf MPH. 2007. SNPSTR: A database of compound microsatellite-SNP markers. Nucleic Acids Res 35: D71–5.

Arandjelovic M, Guschanski K, Schubert G, Harris TR, Thalmann O, Siedel H, Vigilant L. 2009. Two-step multiplex polymerase chain reaction improves the speed and accuracy of genotyping using DNA from noninvasive and museum samples. Mol Ecol Resour 9: 28–36.

Bonatelli IAS, Carstens BC, Moraes EM. 2015. Using next neneration RAD sequencing to isolate multispecies microsatellites for *Pilosocereus* (Cactaceae). PLoS One 10: e0142602.

Buchan JC, Archie EA, Van Horn RC, Moss CJ, Alberts SC. 2005. Locus effects and sources of error in noninvasive genotyping. Mol Ecol Resour 5: 680–683.

Carlson KD, Sudmant PH, Press MO, Eichler EE, Shendure J, Queitsch C. 2015. MIPSTR: a method for multiplex genotyping of germline and somatic STR variation across many individuals. Genome Res 25: 750–761.

Carpenter ML, Buenrostro JD, Valdiosera C, Schroeder H, Allentoft ME, Sikora M, Rasmussen M, Gravel S, Guillén S, Nekhrizov G, et al. 2013. Pulling out the 1%: Whole-Genome capture for the targeted enrichment of ancient dna sequencing libraries. Am J Hum Genet 93: 852–864.

Chikhi R, Medvedev P. 2014. Informed and automated k-mer size selection for genome assembly. Bioinformatics 30: 31–37.

Chiou KL, Bergey CM. 2015. FecalSeq: methylation-based enrichment for noninvasive population genomics from feces. bioRxiv 10.1101/032870.

Ellegren H 2004. Microsatellites: simple sequences with complex evolution. Nat Rev Genet 5: 435–445.

Fan H, Chu J-Y. 2007. A Brief Review of Short Tandem Repeat Mutation. Genomics Proteomics Bioinformatics 5: 7–14.

Fordyce SL, Mogensen HS, Børsting C, Lagacé RE, Chang C-W, Rajagopalan N, Morling N. 2015. Second-generation sequencing of forensic STRs using the Ion Torrent^TM^ HID STR 10-plex and the Ion PGM^TM^. Forensic Sci Int Genet 14: 132–140.

Fu Q, Meyer M, Gao X, Stenzel U, Burbano HA, Kelso J, Pääbo S. 2013. DNA analysis of an early modern human from Tianyuan Cave, China. Proc Natl Acad Sci U S A 110: 2223–2227.

Gnirke A, Melnikov A, Maguire J, Rogov P, LeProust EM, Brockman W, Fennell T, Giannoukos G, Fisher S, Russ C, et al. 2009. Solution hybrid selection with ultra-long oligonucleotides for massively parallel targeted sequencing. Nat Biotech 27: 182–189.

Guichoux E, Lagache L, Wagner S, Chaumeil P, Léger P, Lepais O, Lepoittevin C, Malausa T, Revardel E, Salin F, et al. 2011. Current trends in microsatellite genotyping. Mol Ecol Resour 11: 591–611.

Gymrek M, Golan D, Rosset S, Erlich Y. 2012. lobSTR: A short tandem repeat profiler for personal genomes. Genome Res 22: 1154–1162.

Haak W, Lazaridis I, Patterson N, Rohland N, Mallick S, Llamas B, Brandt G, Nordenfelt S, Harney E, Stewardson K, et al. 2015. Massive migration from the steppe was a source for Indo-European languages in Europe. Nature 522: 207–211.

Harris RS. 2007. Improved pairwise alignment of genomic DNA. PhD dissertation. The Pennsylvania State University.

Hoban SM, Gaggiotti OE, Bertorelle G. 2013. The number of markers and samples needed for detecting bottlenecks under realistic scenarios, with and without recovery: A simulation-based study. Mol Ecol 22: 3444–3450.

Hu X, Yuan J, Shi Y, Lu J, Liu B, Li Z, Chen Y, Mu D, Zhang H, Li N, et al. 2012. pIRS: Profile-based Illumina pair-end reads simulator. Bioinformatics 28: 1533–1535.

Kent WJ. 2002. BLAT—The BLAST-Like Alignment Tool. Genome Res 12: 656–664.

Kistler L, Ratan A, Godfrey LR, Crowley BE, Hughes CE, Lei R, Cui Y, Wood ML, Muldoon KM, Andriamialison H, et al. 2015. Comparative and population mitogenomic analyses of Madagascar’s extinct, giant “subfossil” lemurs. J Hum Evol 79: 45–54.

Koboldt DC, Chen K, Wylie T, Larson DE, McLellan MD, Mardis ER, Weinstock GM, Wilson RK, Ding L. 2009. VarScan: Variant detection in massively parallel sequencing of individual and pooled samples. Bioinformatics 25: 2283–2285.

Lander ES, Waterman MS. 1988. Genomic mapping by fingerprinting random clones: a mathematical analysis. Genomics 2: 231–239.

Li H 2013. Aligning sequence reads, clone sequences and assembly contigs with BWA-MEM. arXiv. http://arxiv.org/abs/1303.3997.

Li H, Durbin R. 2009. Fast and accurate short read alignment with Burrows–Wheeler transform. Bioinformatics 25: 1754–1760.

Li H, Handsaker B, Wysoker A, Fennell T, Ruan J, Homer N, Marth G, Abecasis G, Durbin R. 2009. The Sequence Alignment/Map format and SAMtools. Bioinformatics 25: 2078–2079.

Manichaikul A, Mychaleckyj JC, Rich SS, Daly K, Sale M, Chen WM. 2010. Robust relationship inference in genome-wide association studies. Bioinformatics 26: 2867–2873.

Mckelvey KS, Schwartz MK. 2004. Genetic Errors Associated With Population Estimation Using Non-Invasive Molecular Tagging: Problems and New Solutions. J Wildl Manage 68: 439–448.

Melsted P, Pritchard JK. 2011. Efficient counting of k-mers in DNA sequences using a bloom filter. BMC Bioinformatics 12: 333.

Meyer M, Kircher M. 2010. Illumina sequencing library preparation for highly multiplexed target capture and sequencing. Cold Springs Harb Protoc 10.1101/pdb.prot5448.

Perry GH. 2014. The Promise and Practicality of Population Genomics Research with Endangered Species. Int J Primatol 35: 55–70.

Perry GH, Marioni JC, Melsted P, Gilad Y. 2010. Genomic-scale capture and sequencing of endogenous DNA from feces. Mol Ecol 19: 5332–5344.

Perry GH, Melsted P, Marioni JC, Wang Y, Bainer R, Pickrell JK, Michelini K, Zehr S, Yoder AD, Stephens M, et al. 2012a. Comparative RNA sequencing reveals substantial genetic variation in endangered primates. Genome Res 22: 602–610.

Perry GH, Reeves D, Melsted P, Ratan A, Miller W, Michelini K, Louis EE, Pritchard JK, Mason CE, Gilad Y. 2012b. A Genome Sequence Resource for the Aye-Aye (Daubentonia madagascariensis), a Nocturnal Lemur from Madagascar. Genome Biol Evol 4: 126–135.

Quéméré E, Amelot X, Pierson J, Crouau-Roy B, Chikhi L. 2012. Genetic data suggest a natural prehuman origin of open habitats in northern Madagascar and question the deforestation narrative in this region. Proc Natl Acad Sci USA 109: 13028–13033.

Quinlan AR, Hall IM. 2010. BEDTools: A flexible suite of utilities for comparing genomic features. Bioinformatics 26: 841–842.

Richard AF. 1985. Social boundaries in a Malagasy Prosimian, the Sifaka (Propithecus verreauxi). Int J Primatol 6: 553–568.

Scheible M, Loreille O, Just R, Irwin J. 2014. Short tandem repeat typing on the 454 platform: Strategies and considerations for targeted sequencing of common forensic markers. Forensic Sci Int Genet 12: 107–119.

Schlötterer C, Tautz D. 1992. Slippage synthesis of simple sequence DNA. Nucleic Acids Res 20: 211–215.

Schoebel CN, Brodbeck S, Buehler D, Cornejo C, Gajurel J, Hartikainen H, Keller D, Leys M, Říčanová Š, Segelbacher G, et al. 2013. Lessons learned from microsatellite development for nonmodel organisms using 454 pyrosequencing. J Evol Biol 26: 600–611.

Seguin-Orlando A, Schubert M, Clary J, Stagegaard J, Alberdi MT, Prado JL, Prieto A, Willerslev E, Orlando L. 2013. Ligation bias in illumina next-generation DNA libraries: implications for sequencing ancient genomes. PLoS One 8: e78575.

Seo SB, Ge J, King JL, Budowle B. 2014. Reduction of stutter ratios in short tandem repeat loci typing of low copy number DNA samples. Forensic Sci Int Genet 8: 213–218.

Smith TF, Waterman MS. 1981. Identification of common molecular subsequences. J Mol Biol 147: 195–197.

Snyder-Mackler N, Majoros WH, Yuan ML, Shaver AO, Gordon JB, Kopp GH, Schlebusch SA, Wall JD, Alberts SC, Mukherjee S, et al. 2016. Efficient Genome-Wide Sequencing and Low-Coverage Pedigree Analysis from Noninvasively Collected Samples. Genetics 203: 699–714.

Trapnell C, Pachter L, Salzberg SL. 2009. TopHat: Discovering splice junctions with RNA-Seq. Bioinformatics 25: 1105–1111.

Vartia S, Villanueva-Cañas JL, Finarelli J, Farrell ED, Collins PC, Hughes GM, Carlsson JEL, Gauthier DT, McGinnity P, Cross TF, et al. 2016. A novel method of microsatellite genotyping-by-sequencing using individual combinatorial barcoding. R Soc Open Sci 3: 150565.

Veeramah KR, Hammer MF. 2014. The impact of whole-genome sequencing on the reconstruction of human population history. Nat Rev Genet 15: 149–162.

Willems T, Gymrek M, Highnam G, Mittelman D, Erlich Y. 2014. The landscape of human STR variation. Genome Res 24: 1894–1904.

